# Genome size drives morphological evolution in organ-specific ways

**DOI:** 10.1101/2021.08.17.456171

**Authors:** Michael W. Itgen, Dustin S. Siegel, Stanley K. Sessions, Rachel Lockridge Mueller

## Abstract

Morphogenesis is an emergent property of biochemical and cellular interactions during development. Genome size and the correlated trait of cell size can influence these interactions through its effects on developmental rate and tissue geometry, ultimately driving the evolution of morphology. We tested the effects of genome size and body size evolution on heart and liver morphology using nine species of the salamander genus *Plethodon* (genome sizes 29.3–67 Gb). Our results show that whole organ size is determined by body size, whereas tissue structure changes dramatically with evolutionary increases in genome size. In the heart, increased genome size is correlated with a reduction of myocardia in the ventricle, yielding proportionally less force–producing mass and more empty space. In the liver, increased genome size is correlated with fewer and larger vascular structures, positioning hepatocytes farther from the circulatory vessels that transport key metabolites. Although these structural changes should have obvious impacts on organ function, their effects on organismal performance and fitness are likely negligible because low metabolic rates in salamanders relax selective pressure on key metabolic organ performance. Overall, this study reveals the effects of large genome and cell size on the developmental systems producing the heart and liver.

## Introduction

The evolutionary trajectories of morphological traits reflect the interaction between possible phenotypes for a species and selection to match these phenotypes to the environment. While optimal trait values theoretically exist for any species in any environment, in reality, there are evolutionary constraints that define a limited set of possible phenotypes from all theoretical ones across morphospace (Alberch, 1982). Evolutionary constraints can exist at all levels of biology and can involve both developmental and physical limitations that can introduce bias or prevent certain phenotypes from evolving (Maynard Smith et al., 1985; Arnold, 1992; Brakefield, 2006; Gerber, 2014).

The separation of possible from theoretical phenotypes in morphospace is often defined by whether the phenotypes can be produced by the species’ developmental system (Alberch, 1982; Salazar-Ciudad, 2006). The process of development is an emergent property of biochemical and cellular interactions that direct morphogenesis. Gene expression and signal transduction networks instruct populations of cells to divide, differentiate, migrate, and coalesce (Alberch, 1982; Oster et al., 1988). In addition, cells influence their neighbors, relaying positional and deterministic information to one another that results in the induction of tissue formation and organogenesis. Molecular- and cell-level processes are both required; genes and gene products alone cannot produce tissues and organs without the organizational level of cells, and cells cannot organize without molecular instruction. When genes, proteins, or cells evolve within this system, any new variant can potentially alter these collective interactions and change the outcome of development.

Alberch (1982) proposed that the primary forces underlying the evolution of morphology are changes to the biochemical and cellular interactions involved in development. Overall, development is robust to change, and variation — both mutational and epigenetic — can often have no impact on the resulting phenotype (Lewontin, 1972; Oster and Alberch, 1982; Wagner, 2011; Uller et al., 2018). Thus, many possible interactions between molecules and cells exist that will yield the same developmental outcome (Alberch, 1982; Oster and Alberch, 1982). However, some such changes will result in the developmental system producing a different phenotype (Oster and Alberch, 1982; Uller et al., 2018). These changes cross a theoretical threshold called a bifurcation boundary, which bounds the amount of variation permittable within a developmental system that still produces the same outcome (Oster and Alberch, 1982). This variation can exist in many parameters, including: sequence, structure, function, and interaction of genes and proteins; rates of diffusion and biochemical reactions; morphology and motility of cells; rates of cellular division and differentiation; and the size and organization of tissues and structures (Alberch, 1983; Brakefield et al., 2003; Mallarino et al., 2011; Keyte and Smith, 2014; Powder et al, 2015). Variations in these parameters can collectively alter morphogenesis and result in larger scale changes that impact the rate of development, the sequential timing of developmental events, or pattern formation, pushing the system across a bifurcation boundary (Oster and Alberch, 1982).

Genome size is a trait that can directly impact developmental systems through its effects on cell biology (Gregory, 2005). There is a strong positive correlation between genome size and cell size, resulting in cells becoming larger as DNA accumulates (Gregory, 2005; D’Ario et al., 2021). Large cell size has clear impacts on cell morphology, shifting the ratio between surface area and volume and causing the scaling of intracellular organelles (Marshall et al., 2012). Genome and cell size together have been shown to impact developmental systems through two mechanisms (Gregory, 2005). First, large genome and cell sizes slow developmental rate by causing longer cell cycles and slower rates of cell migration and differentiation (Sessions and Larson, 1987; Schmidt and Roth, 1993; Vinogradov, 1999). This slowing can ultimately shift the timing of developmental events (i.e., heterochrony) and impact the dynamics of morphogenesis

(Alberch et al., 1979; Gould, 1985). Changes in cell size also impact the final outcomes of development: the structures of tissues and organs (Alberch and Alberch, 1981; Hanken and Wake, 1993; Roth et al., 1993). Tissues and organs are a function of both cell size and cell number. Therefore, evolutionary increases in genome and cell size can result in organs and organisms that are composed of fewer, larger cells, assuming body size and organ size remain constant (Hanken and Wake, 1993). Morphogenesis must then occur with fewer cells, and the final organ or structure must be able to maintain function with fewer, larger cells.

Empirical studies of morphological evolution have suggested that the impacts of increased genome and cell size — mediated through alterations to the developmental system — can vary extensively across different organs, structures, and species (Fankhauser, 1945; Hanken and Wake, 1993; Roth et al., 1993; Snyder and Sheafor, 1999; Womack et al., 2019). Amphibians, particularly salamanders, have provided a powerful system for studying these patterns and processes due to their enormous range in genome and cell size (Gregory, 2005; Sessions, 2008; Sessions and Wake, 2021). Much of this work has focused on the brain and nervous system, the skeletal system, and the circulatory system. Within the salamander brain, dramatic changes in gross morphology and tissue organization are connected to larger genome and cell sizes as well as reductions in cell numbers (Roth et al., 1993; Roth and Walkowiak, 2015). Several brain regions including the thalamus, praetectum, and midbrain become proportionally larger primarily through a reduction in the forebrain (Roth et al., 1988, 1990). Larger and fewer cells also impact tissue organization, leading to an increase in gray matter relative to white matter as well as increased cell density in several tissues (Roth et al., 1990). In addition, slower rates of cell proliferation and migration caused by large genome and cell size results in a decrease in lamination within the tectum mesencephali (Schmidt and Roth, 1993).

Amphibian skeletal elements also show a variety of effects because of genome and cell size increase (Alberch and Alberch, 1981; Hanken and Wake, 1993; Womack et al., 2019). In tropical lungless salamanders with exceptionally large genomes, slower development rates appear to be connected to paedomorphic morphologies including fenestrated skulls, highly reduced or even absent phalangeal elements in the digits, and extensively webbed feet (Wake, 1966; Alberch and Alberch, 1981; Alberch, 1983; Jaekel and Wake, 2007; Decena-Segarra et al., 2020). In addition, the relationship between cell size and body size — also known as biological size (Hanken and Wake, 1993) — appears to disrupt skeletal development when genome size increases (and/or body size decreases) because the prerequisite tissues that form the bones are reduced to significantly fewer cells (Hanken, 1982, 1984; Wake, 1991). In some cases, the carpal and tarsal elements of the feet remain cartilaginous and are often fused due to a failure to separate during development (Wake, 1966; Alberch and Alberch, 1981). In extreme cases, abnormal morphologies and losses of skeletal elements occur, often asymmetrically within single individuals. The same skeletal elements can show different patterns of change accompanying increased cell size depending on body size (Wake, 1991; Hanken and Wake, 1993). Comparative analyses have found evidence that miniaturized salamanders have undergone genome size reduction because of the consequences of limited cell number as a developmental constraint (Decena-Segarra et al., 2020).

Large genome and cell size has also impacted the circulatory and excretory systems. Red blood cell size is positively correlated with genome size across vertebrates (Gregory, 2001). Evolutionary increases in red blood cell size are accompanied by wider capillary diameters (Snyder and Sheafor, 1999). Multiple lineages of miniaturized salamanders have secondarily evolved enucleated red blood cells to reduce cell size and permit passage through small capillaries (Villalobos et al., 1988; Mueller et al., 2008; Itgen et al., 2019; Decena-Segarra et al., 2020). In contrast, experimental induction of polyploidy (and thus increased cell size) did not result in proportional increases in the diameter of pronephric tubules, which are excretory structures similar in morphology to capillaries; however, increased ploidy did alter the morphology and number of cells comprising the structure (Fankhauser, 1945).

The diversity of morphological outcomes in the studies carried out to date suggests that the fundamental rules governing the effects of genome and cell size on morphological evolution will be revealed through the analysis of additional organs and the synthesis of results across species and organ systems. As a step towards this goal, our study investigates how increases in genome and cell size impact the morphology of the heart and liver. These two previously unexplored organs differ in morphogenesis and function; the heart has a kinetic biomechanical function and the liver has a biochemical and secretory function. Our study system is the lungless salamander genus *Plethodon*, which includes species with dramatic differences in genome size (23 – 67 Gb) and cell size, but uniformity or lower diversity in potentially confounding variables that could also impact organ structure and function (life history, body size; Highton, 1995; Newman et al., 2016). Using diffusible iodine-based contrast-enhanced computed tomography (diceCT) and histology applied to nine focal *Plethodon* species, we quantified body size, organ size, and several measures of tissue composition and geometry: 1) proportion of the heart wall comprised of cardiomyocytes versus lumen, 2) proportion of the liver comprised of hepatocytes versus vascular openings, and 3) the numbers and sizes of distinct vascular structures (sinusoids, veins, arteries) in the liver. We used phylogenetic comparative methods to test whether these variables correlate with evolutionary changes in genome and cell size. Based on our findings, we propose hypotheses connecting cell size, developmental system perturbation, and morphology. More generally, we discuss how relaxed functional constraints on organ performance can allow the evolution of a range of structurally different, but functionally equivalent, morphologies.

## Methods

### Animal collection

We collected five adult individuals of *P. cinereus, P. cylindraceus, P. dunni, P. glutinosus, P. idahoensis, P. metcalfi, P. montanus, P. vandykei*, and *P. vehiculum*. Permit IDs and locality data can be found in the supplemental data. Salamanders were collected and euthanized in buffered 1% MS-222, fixed in buffered formalin, and transferred through a graded series of ethanol (10%, 30%, 50%, 70%) before storage in 70% ethanol indefinitely. The protocols for animal research, husbandry, and euthanasia were approved by the Institutional Animal Care and Use Committee of Colorado State University (17-7189A).

### diceCT data generation and processing

We used diffusible iodine-based contrast-enhanced computed tomography (diceCT) to collect data on liver and heart volumes (Gignac et al., 2016). In order to produce the diceCT scans, specimens were stained in a 1% I_2_KI solution for 2 days. Specimens were scanned twice using a Bruker SkyScan 1173 at the Karel F. Liem Bioimaging Center, Friday Harbor Laboratories, University of Washington. Scans were set to 85 kV and 90 uA with a 1 mm aluminum filter to reduce beam hardening. We first produced full body scans to measure liver volumes at a resolution ranging from 14.9–17 μm, depending on the size of the specimens. We then produced higher resolution scans for the heart, which were scanned at a resolution ranging from 7.1–9.9 μm. After the specimens were scanned, the I_2_KI was rinsed out using several changes of 70% ethanol and the specimens were stored in 70% ethanol. All of the specimens were treated uniformly to standardize for any I_2_KI-related artefacts or tissue shrinkage that might impact the subsequent histological analyses.

diceCT scans were reconstructed using NRecon (Bruker, 2005–2011) following standard operating procedures including optimal x/y alignment, ring artifact reduction, beam hardening correction, and a post-alignment. Data visualization and analysis were accomplished using 3DSlicer (Fedorov et al., 2012). Liver and heart ventricle volumes were calculated through segmentation of the liver and heart from each individual.

### Organ measurement

Snout–vent lengths were measured to the nearest 0.01 mm for each individual using digital calipers. The hearts and livers were then dissected out from the specimens and were embedded in plastic following standard protocols (Humason, 1962). Tissues were sectioned at 4 µm and stained with hematoxylin for 4 minutes and toluidine for 3 minutes. Sections were mounted and then visualized using a compound microscope. Images used in the analysis were minimally edited to remove blood cells that obstructed vascular structures or the lumen of the ventricle. Five images were taken from unique slides at 20x magnification for each individual and a mean value was calculated for each morphological trait per individual. Each image was converted to greyscale and a thresholding method was used to collect the morphometric data. ImageJ was used for all image processing and analysis (Schneider et al., 2012).

Amphibian livers are primarily comprised of hepatic tissue that is permeated by the circulatory system (Akiyoshi and Inoue, 2012). Liver circulatory structures consist of hepatic arteries that provide oxygen, portal veins that bring nutrients and toxins to the liver, and sinusoids, which are specialized capillaries where oxygen-rich blood from hepatic arteries and nutrient-rich blood from portal veins mix (Elias and Bengelsdorf, 1952). Liver tissue is generally arranged into many hepatic lobules that are centered by portal triads – an arrangement of hepatic arteries, portal veins, and bile ducts (Elias and Bengelsdorf, 1952). The network of sinusoids gives the hepatic tissue a cord-like appearance in most vertebrate taxa, with hepatocytes forming cords that are 1–2 cells thick; this morphology increases the surface area of each hepatocyte that is in contact with circulating blood (Elias and Bengelsdorf, 1952). However, some species of salamanders have a many-cell thick arrangement of hepatic cords (Akioyshi and Inoue, 2012). For the liver, we measured the total area of each histological section that was comprised of tissue (primarily hepatocytes) versus vascular openings and the number and size of distinct vascular structures (sinusoids, veins, arteries) (Fig. S1). Twenty nuclei and cells were also measured for each individual to collect data on hepatocyte nuclear and cell area.

Amphibians have a single, thin-walled ventricle that has a central chamber surrounded by a highly trabeculated network of myocardium, which is a characteristic of ectotherms (Stephenson et al., 2017). Heart morphology in *Plethodon* also reflects the lack of lungs in the family Plethodontidae, which has been accompanied by a loss of complete atrial septation (Lewis and Hanken, 2017). For the heart ventricles, we measured the amount of myocardial area in the ventricle walls versus lumen (Fig. S1). We focused on this because the trabeculated myocardium makes it difficult to define the edges of the ventricle chamber. We did not measure any characteristics of the atria because they lack distinct internal structure and their elastic nature made accurate volumetric measurements impossible.

### Genome size measurement

Genome size was measured using the Feulgen-staining method on fixed erythrocytes following the protocol of Sessions and Larson (1987). *Ambystoma mexicanum* (32 Gb) was used as a standard to calculate the genome sizes of the other species. The *A. mexicanum* were acquired from the Ambystoma Genetic Stock Center at the University of Kentucky. Erythrocytes were extracted from the *Plethodon* and *Ambystoma* specimens fixed in buffered formalin and transferred to microscope slides to produce blood smears. We collected blood smears from 3–5 individuals per species. The cells were hydrated for 3 minutes in distilled water, permeabilized in 5 N HCl for 20 minutes at 20°C, and then rinsed three times in distilled water. Nuclei were stained with Schiff’s reagent for 90 minutes at 20°C, destained in 0.5% sodium metabisulfite three times for 5 minutes each, and then rinsed in distilled water three times. The stained cells were dehydrated in a graded series of 70%, 95%, and 100% ethanol, dried, and mounted. We photographed 2–12 (x□ = 5.5) nuclei per individual under 100x, and the integrated optic densities (IOD) were measured using IMAGE PRO software (Media Cybernetics, Rockville, Maryland, USA). Genome sizes were calculated by comparing the average IODs of the experimental species to the IOD of the standard. Nuclear areas were also measured for each erythrocyte using the IMAGE PRO software.

### Phylogeny

We estimated the phylogenetic relationships among the 9 species of *Plethodon* used in this study to account for phylogenetic non-independence in our analyses. DNA sequences for the mtDNA gene *cytb* and the nuclear gene *Rag1* were obtained from NCBI (https://www.ncbi.nlm.nih.gov/genbank; Table S1). The *cytb* and *Rag1* datasets were aligned independently using MUSCLE in MEGA v7 with default parameters and trimmed to 629 and 1,467 basepairs, respectively (Edgar, 2004; Kumar et al., 2016). We selected a GTR+G+I nucleotide substitution model by running PartitionFinder 2 on each codon position for both genes (Lanfear et al., 2017). The phylogeny was estimated using Bayesian inference with MrBayes v3.2.5 (Ronquist et al., 2012). The analysis ran with four chains (3 heated, 1 cold) for 10 million generations with sampling occurring every 1,000 and the first 10% of the sampled trees discarded as burnin.

### Data analysis

We first tested if organ morphological traits, genome size, and SVL were phylogenetically dependent by estimating phylogenetic signal using Blomberg’s K based on our midpoint-rooted phylogeny. Blomberg’s K was calculated using the *phytools* package implemented in R v 3.4.2 (Revell, 2012; R core team, 2016). We then log-transformed all variables to account for non-normal distributions and calculated phylogenetic independent contrasts (PICs) for each variable using the R package *phytools* (Revell, 2012). We ran a MANOVA and subsequent post-hoc univariate ANOVA analysis on a model including SVL and genome size as predictor variables and all morphological data as response variables to test if organ morphology is correlated with genome size and/or SVL while accounting for phylogeny. Our response variables included liver size, ventricle size, total area comprised of muscle in the ventricle, number of vascular structures in the liver, average size of the vascular structures in the liver, and the total area comprised of hepatic tissues in the liver. The MANOVA and univariate ANOVA analyses were conducted using R v 3.4.2 (R core team, 2016).

We applied a Brownian motion model of evolution to all variables because there is evidence that genome size evolves by largely stochastic processes in direct-developing salamanders such as *Plethodon* (Mueller et al., preprint). Because we hypothesize that organ morphology is shaped by the evolution of genome and cell size, we predict that the heart and liver morphological traits are not evolving towards adaptive optima. We visualized the magnitude and direction of changes in genome size across the 9 species of *Plethodon* in the study using the contMap function in the R package *phytools* on the estimated topology (Revell, 2012).

## Results

The genome size estimates for the 9 *Plethodon* species ranged from 29.3–67.0 Gb (Figure 1; Table 1). The genome sizes estimated in this study were generally larger than those previously published or fell within the higher range of published estimates (Gregory, 2021). We estimated larger genome sizes for *P. cinereus* (29.3 Gb vs. 25.6 Gb), *P. dunni* (52.3 Gb vs. 46.5 Gb), and *P. vehiculum* (46.4 Gb vs. 39.1 Gb). The genome size estimate for *P. glutinosus* was comparable to the higher published estimates for the species (38.9 Gb vs. 42.1 Gb), but is much larger than the average published value for this species (X = 28.0 Gb). We also estimated that *P. idahoensis* has the largest genome size in the genus at 67.0 Gb (Sessions, 2008; Newman et al., 2016; Gregory, 2021). Conversely, we estimated a significantly smaller genome size for *P. vandykei* (54.6 Gb vs. 67.8 Gb), which was previously considered to have the largest genome size in the genus. Intraspecific variation in genome size measurements could, in principle, reflect true variation within and among populations, changes in taxonomic assignments, and/or technical discrepancies across studies. We circumvented these sources of uncertainty by collecting our own genome size data on the same organisms we used for morphological analysis. The areas of nuclei from both hepatocytes and erythrocytes are highly correlated with these genome size measurements, which supports the accuracy of these data (Fig. 4; Table 1).

**Table 1.**
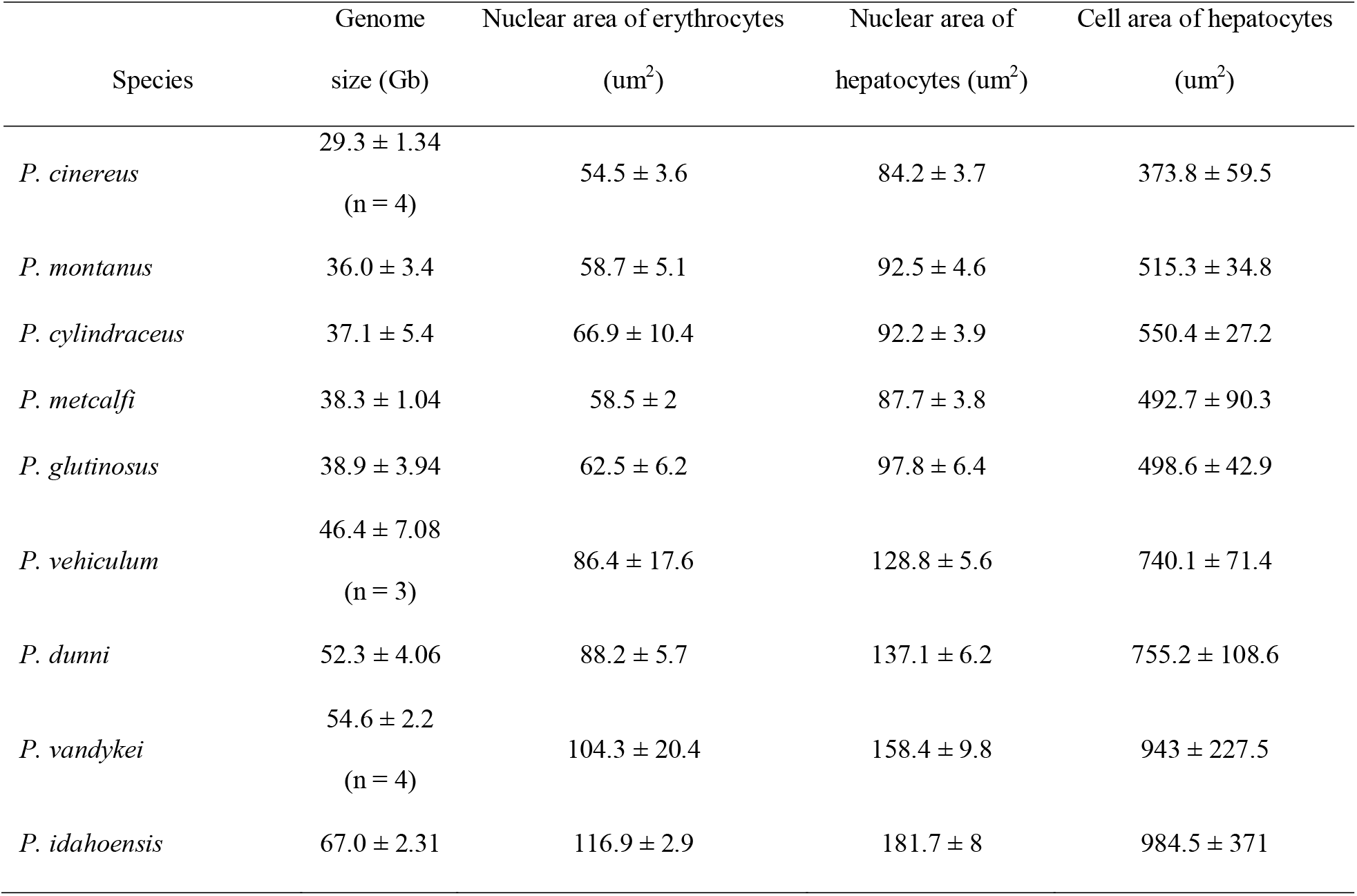
Mean and standard deviation for measurements of genome size, nuclear areas of erythrocytes and hepatocytes, and cell areas of hepatocytes. n = 5 individuals unless otherwise noted; 3–5 erythrocyte nuclei and 20 hepatic nuclei and cells were measured per individual.

| Species | Genome size (Gb) | Nuclear area of erythrocytes (um<sup>2</sup>) | Nuclear area of hepatocytes (um<sup>2</sup>) | Cell area of hepatocytes (um<sup>2</sup>) |
| --- | --- | --- | --- | --- |
| *P. cinereus* | 29.3 ± 1.34<br>(n = 4) | 54.5 ± 3.6 | 84.2 ± 3.7 | 373.8 ± 59.5 |
| *P. montanus* | 36.0 ± 3.4 | 58.7 ± 5.1 | 92.5 ± 4.6 | 515.3 ± 34.8 |
| *P. cylindraceus* | 37.1 ± 5.4 | 66.9 ± 10.4 | 92.2 ± 3.9 | 550.4 ± 27.2 |
| *P. metcalfi* | 38.3 ± 1.04 | 58.5 ± 2 | 87.7 ± 3.8 | 492.7 ± 90.3 |
| *P. glutinosus* | 38.9 ± 3.94 | 62.5 ± 6.2 | 97.8 ± 6.4 | 498.6 ± 42.9 |
| *P. vehiculum* | 46.4 ± 7.08<br>(n = 3) | 86.4 ± 17.6 | 128.8 ± 5.6 | 740.1 ± 71.4 |
| *P. dunni* | 52.3 ± 4.06 | 88.2 ± 5.7 | 137.1 ± 6.2 | 755.2 ± 108.6 |
| *P. vandykei* | 54.6 ± 2.2<br>(n = 4) | 104.3 ± 20.4 | 158.4 ± 9.8 | 943 ± 227.5 |
| *P. idahoensis* | 67.0 ± 2.31 | 116.9 ± 2.9 | 181.7 ± 8 | 984.5 ± 371 |

**Figure 1.**
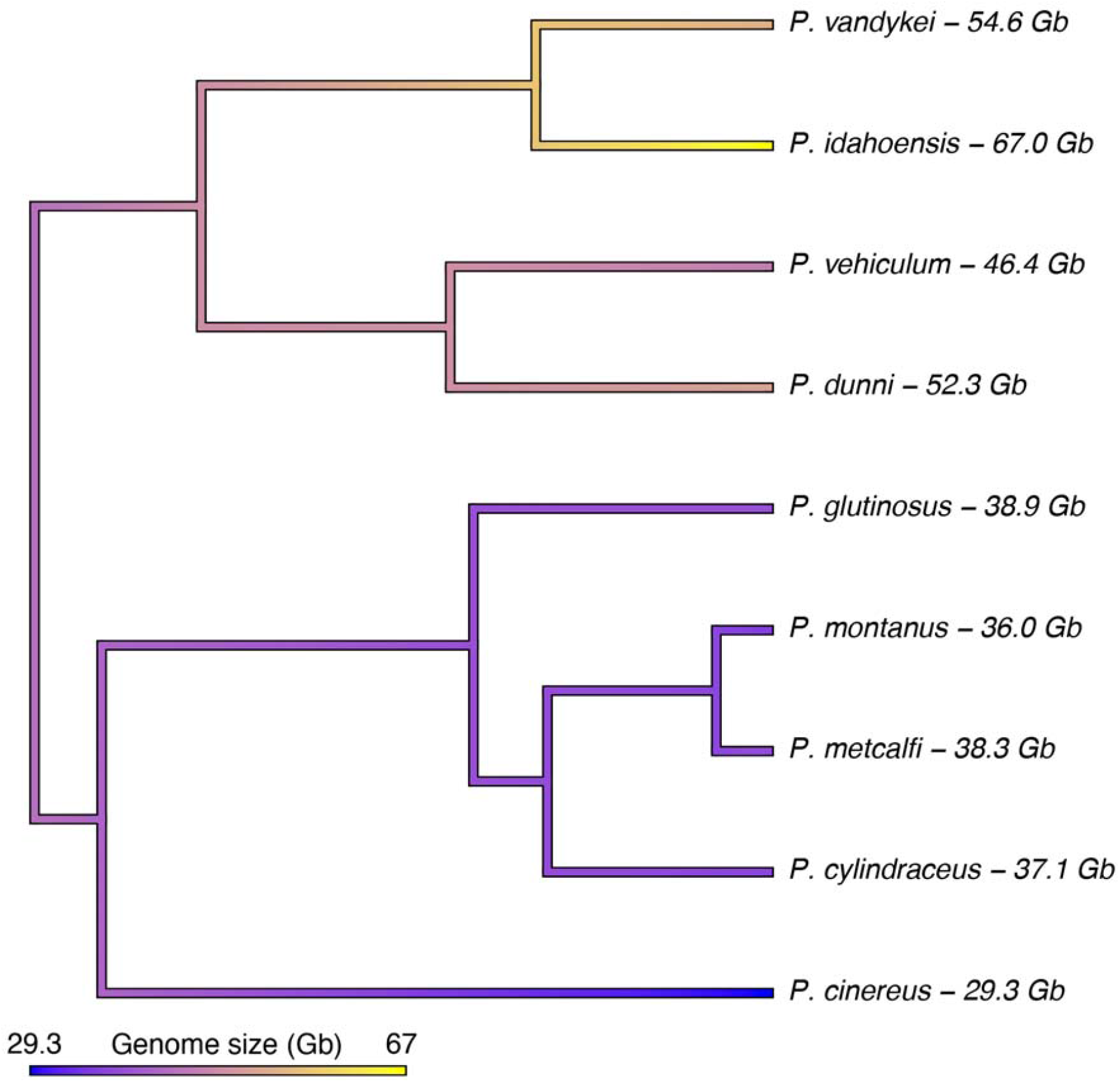
Genome size mapped as a continuous trait onto the topology for the 9 species of *Plethodon* to visualize differences among species.

Morphological and histological trait data are summarized in Table 2. Mean SVL ranged from 40.1 mm in *P. cinereus* to 68.7 mm in *P. dunni*. Mean ventricle volume spanned an order of magnitude across this range of body sizes — from 0.52 mm^3^ in *P. cinereus* to 5.72 mm^3^ in *P. dunni*. Mean liver volume showed a ∼7-fold range across these body sizes, from 15.9 mm^3^ in *P. cinereus* to 112.1 mm^3^ in *P. dunni*. Mean myocardial density showed a ∼3-fold range across species, from 0.04 mm^2^ / section in *P. idahoensis* to 0.118 mm^2^ / section in *P. cinereus*. The mean number of vascular structures in the liver showed a ∼2.5-fold range, from 23.5 / section in *P. idahoensis* to 60.0 in *P. cinereus*. Mean size of vascular structures showed a ∼3-fold range, from 0.0021 mm^2^ in *P. cylindraceus* and *P. glutinosus* to 0.0061 mm^2^ in *P. idahoensis*. Mean hepatic tissue area showed the smallest range across species, from 0.127 mm^2^ in *P. glutinosus* to 0.143 mm^2^ in *P. idahoensis*.

**Table 2.** Mean and standard deviation for genome size and morphological traits. n = 5 individuals unless otherwise noted; 5 slides were measured per individual.

| Species | Snout-vent | Heart |  | Liver |  |  | Mean size of<br>vascular<br>structures (mm<sup>2</sup>) |
| --- | --- | --- | --- | --- | --- | --- | --- |
|  | length | Ventricle |  | Liver |  |  |  |
|  | (mm) | volume | Myocardial | volume | Hepatic tissue | Number of |  |
|  |  | (mm<sup>3</sup>) | area (mm<sup>2</sup>) | (mm<sup>3</sup>) | area (mm<sup>2</sup>) | sinusoids |  |
| *P. cinereus* | 40.1 ± 0.7 | 0.52 ± 0.06 | 0.118 ± 0.01 | 15.9 ± 3.8 | 0.136 ± 0.01 | 60 ± 7.1 | 0.0023 ± 0.0002 |
| *P. montanus* | 49.7 ± 1.1 | 1.44 ± 0.35 | 0.101 ± 0.004 | 33.1 ± 15.9 | 0.14 ± 0.01 | 57.1 ± 4.7 | 0.0024 ± 0.0005 |
| *P. cylindraceus* | 70.4 ± 1.6 | 3.77 ± 1.24 | 0.107 ± 0.01 | 120 ± 72.5 | 0.123 ± 0.01 | 58.4 ± 8.8 | 0.0021 ± 0.0002 |
| *P. metcalfi* | 61.9 ± 4.2 | 3.74 ± 1.63 | 0.106 ± 0.004 | 70.6 ± 41.3 | 0.136 ± 0.01 | 54.7 ± 6.6 | 0.0025 ± 0.0002 |
| *P. glutinosus* | 66.9 ± 2.4 | 3.71 ± 1.63 | 0.101 ± 0.002<br>(n = 4) | 77.9 ± 40.3 | 0.127 ± 0.01 | 59.6 ± 3.5 | 0.0021 ± 0.0002 |
| *P. vehiculum* | 50.5 ± 0.7 | 0.9 ± 0.1 | 0.09 ± 0.004 | 40 ± 5.6 | 0.141 ± 0.003 | 34.4 ± 3.0 | 0.0041 ± 0.0005 |
| *P. dunni* | 68.7 ± 1.2 | 5.72 ± 1.59 | 0.065 ± 0.01 | 112.1 ± 63.1<br>(n = 3) | 0.136 ± 0.004 | 32.4 ± 4.9 | 0.0042 ± 0.0002 |
| *P. vandykei* | 54.2 ± 2.0 | 1.51 ± 0.42 | 0.073 ± 0.01 | 62.2 ± 17.9<br>(n = 4) | 0.142 ± 0.002 | 26.3 ± 4.8 | 0.0054 ± 0.0006 |
| *P. idahoensis* | 58.1 ± 2.1 | 2.63 ± 0.78 | 0.043 ± 0.01 | 40.1 ± 14.7<br>(n = 4) | 0.143 ± 0.004 | 23.5 ± 3.0 | 0.0061 ± 0.0002 |

There was significant phylogenetic signal in genome size (Blomberg’s K = 1.51; *P* = 0.001), myocardial area (Blomberg’s K = 0.97; *P* = 0.019), the number of vascular structures (Blomberg’s K = 2.69; *P* = 0.002), and the average size of vascular structures (Blomberg’s K = 2.6; *P* = 0.002) (Table S2). The results from the MANOVA analysis are presented in Table 3. Body size (SVL) was positively correlated with ventricle size (P = 0.002) and liver size (P < 0.001; Fig. 5, Fig. S2–3). In the heart, genome size was negatively correlated with myocardial area in the ventricle (*P* < 0.001; Fig. 2,6). In the liver, genome size showed both positive and negative correlations with different traits: genome size was positively correlated with the average size of the vascular structures (P < 0.001; Fig 3,6) but negatively correlated with the number of vascular structures (P < 0.001; Fig 3,6). In addition, genome size was positively correlated with the total hepatic tissue area (P = 0.013; Fig. 3,6). There were also significant positive correlations between body size (SVL) and the total hepatic tissue area (P = 0.003) and the average vascular structure size (P = 0.006) in the liver.

**Table 3.**
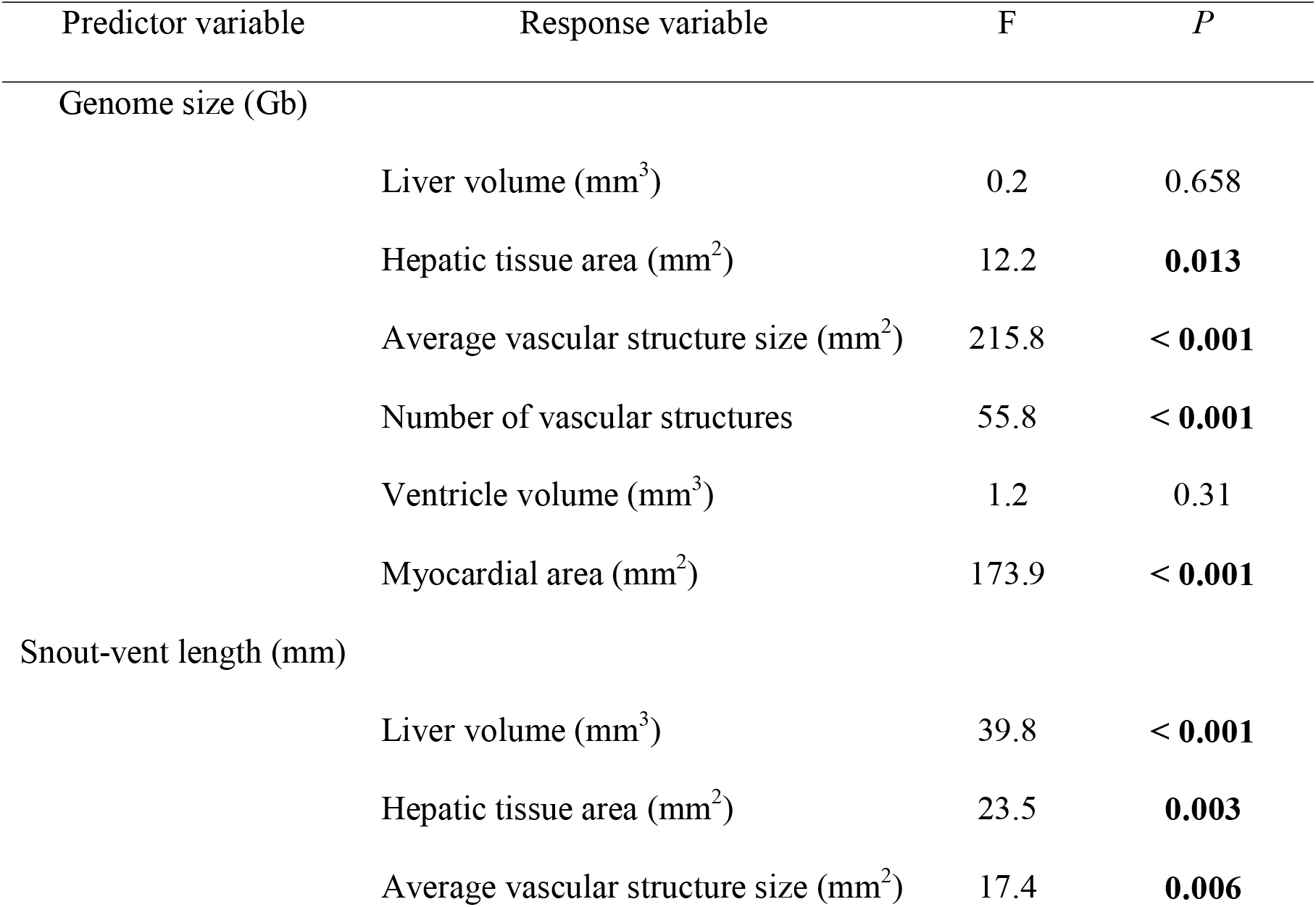
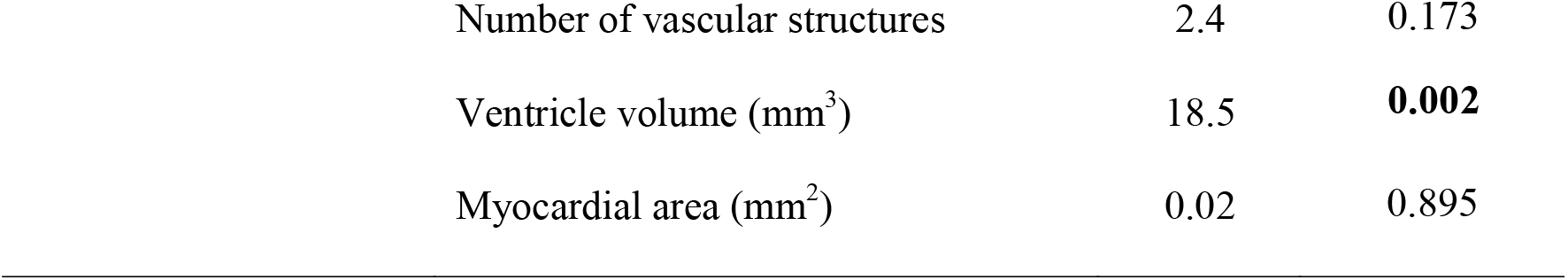
Results from the univariate ANOVA showing the individual interactions between the predictor and response variables.

**Figure 2.**
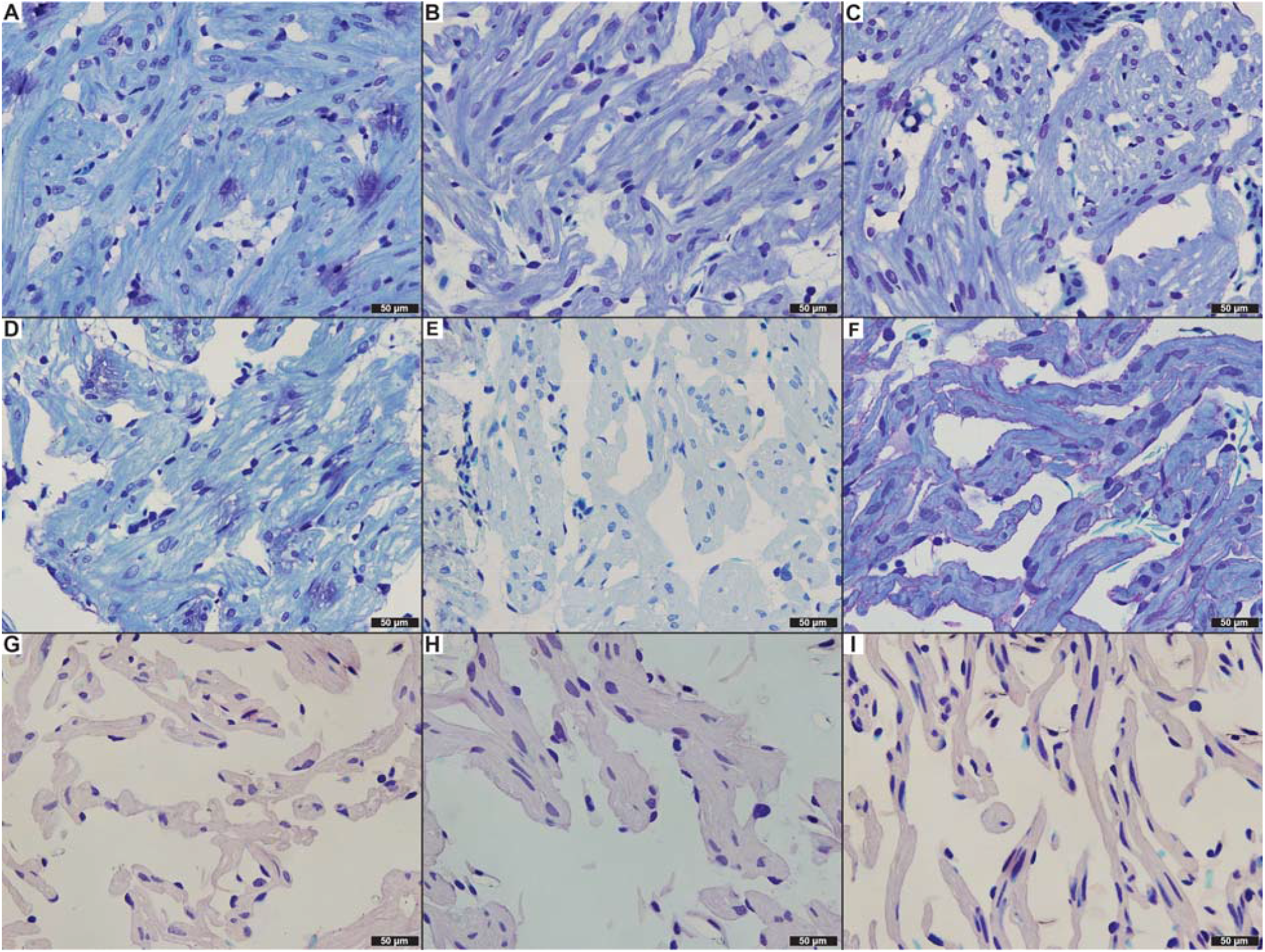
Histological sections of the ventricle at 20x magnification for each species arranged by increasing genome size. A) *P. cinereus*, 29.3 Gb; B) *P. montanus*, 36 Gb; C) *P. cylindraceus*, 37.1 Gb; D) *P. metcalfi*, 38.3 Gb; E) *P. glutinosus*, 38.9 Gb; F) *P. vehiculum*, 46.4 Gb; G) *P. dunni*, 52.3 Gb; H) *P. vandykei*, 54.6 Gb; I) *P. idahoensis*, 67 Gb.

**Figure 3.**
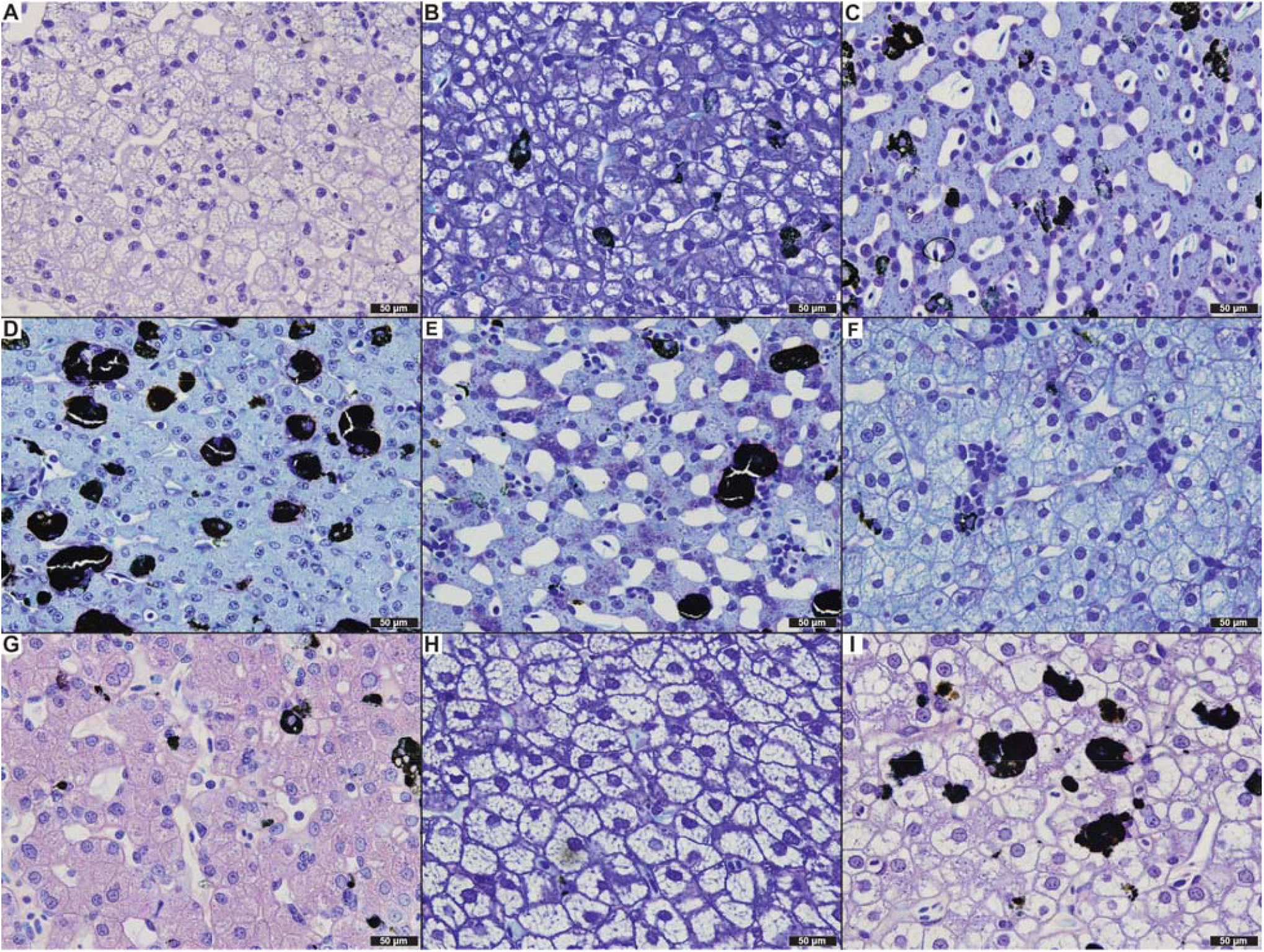
Histological sections of the liver at 20x magnification for each species arranged by increasing genome size. A) *P. cinereus*, 29.3 Gb; B) *P. montanus*, 36 Gb; C) *P. cylindraceus*, 37.1 Gb; D) *P. metcalfi*, 38.3 Gb; E) *P. glutinosus*, 38.9 Gb; F) *P. vehiculum*, 46.4 Gb; G) *P. dunni*, 52.3 Gb; H) *P. vandykei*, 54.6 Gb; I) *P. idahoensis*, 67 Gb.

**Figure 4.**
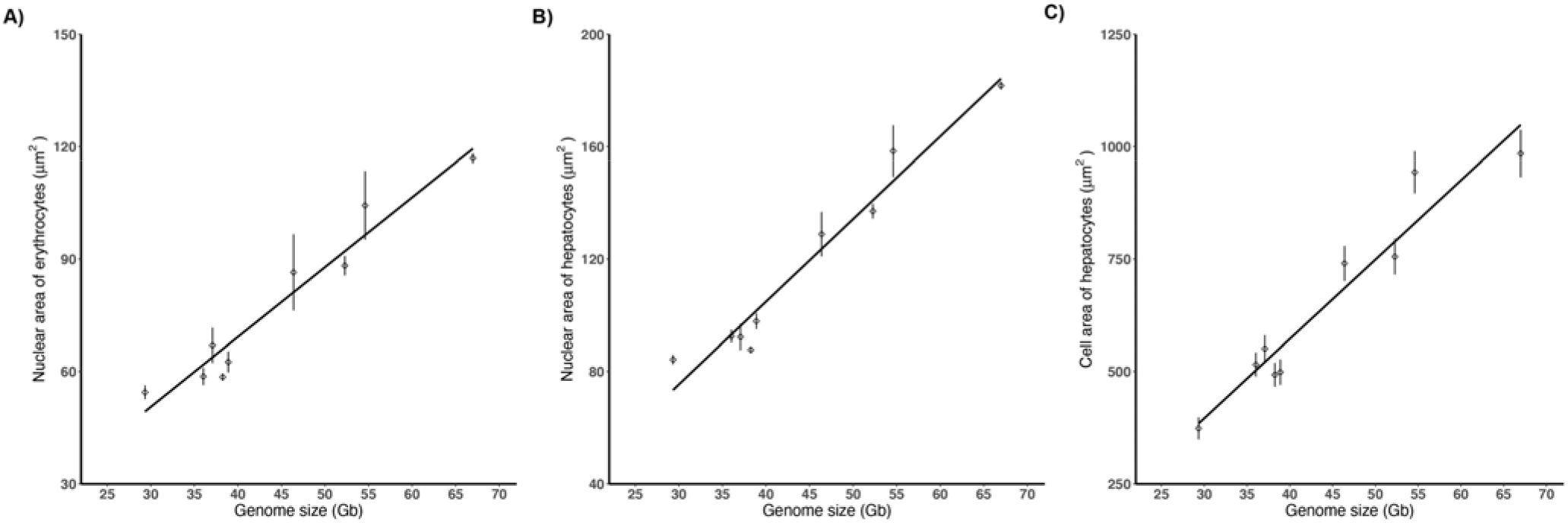
Genome size has a strong positive correlation with the A) nuclear area of erythrocytes, B) nuclear area of hepatocytes, and C) cell area of hepatocytes across the nine species. Error bars represent standard error.

**Figure 5.**
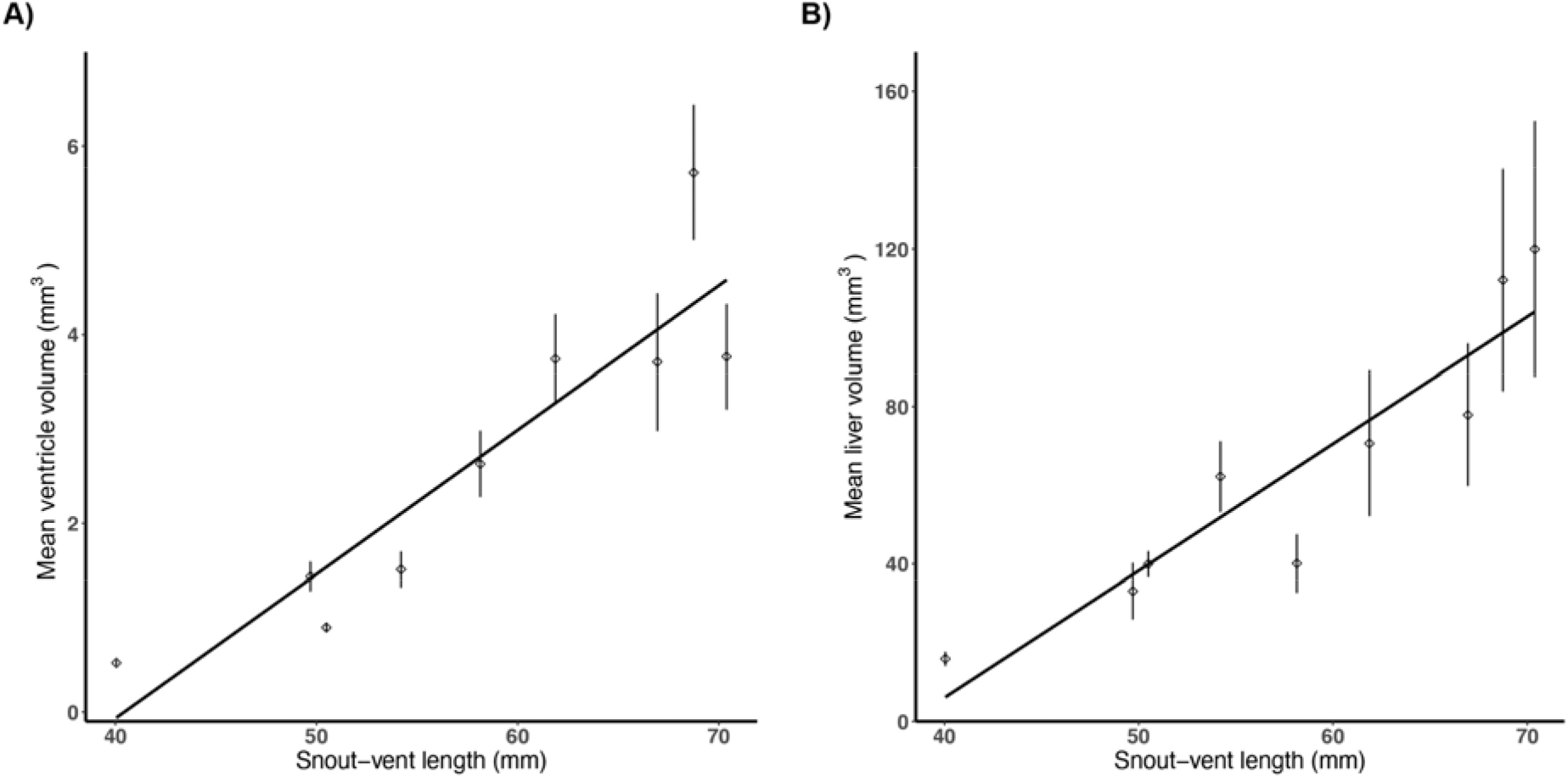
Body size (SVL) is positively correlated with A) ventricle size and B) liver size. Error bars represent standard error.

**Figure 6.**
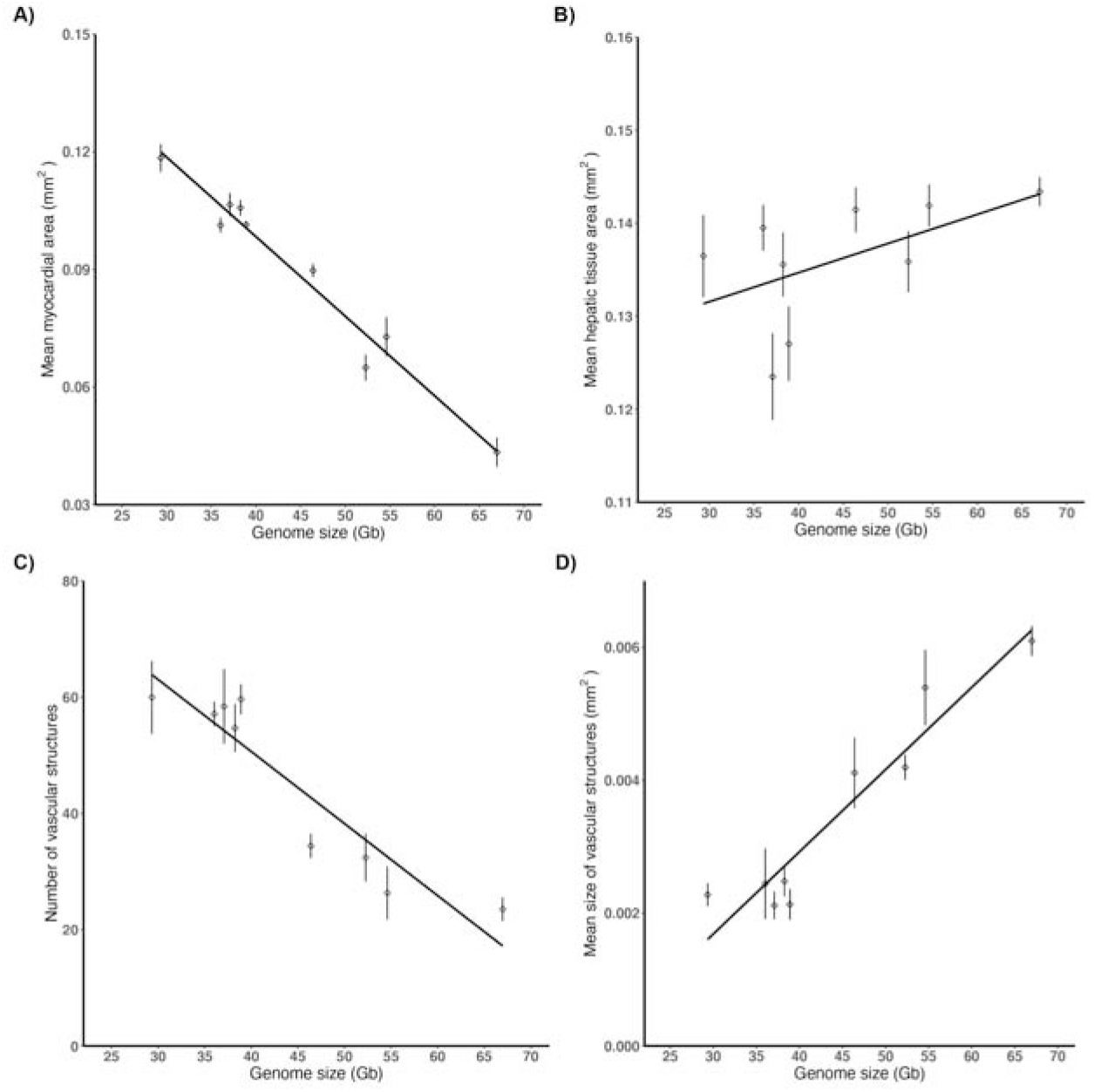
An increase in genome size is correlated with A) a decrease in myocardial area, B) an increase in hepatic tissue area, C) a decrease in the number of vascular structures, and D) an increase the size of vascular structures. Error bars represent standard error.

Hepatic tissue area and average size of the vascular structures in the liver were significantly correlated with both genome size and SVL, suggesting that these traits are determined by the biological size index (BSI) – a relative measure of the total number of cells comprising an organism that is based on organism size and cell size. BSI is calculated by dividing the SVL by the square-root of genome size. We calculated the BSI and found that it is significantly correlated with both of these traits (Fig. 7).

**Figure 7.**
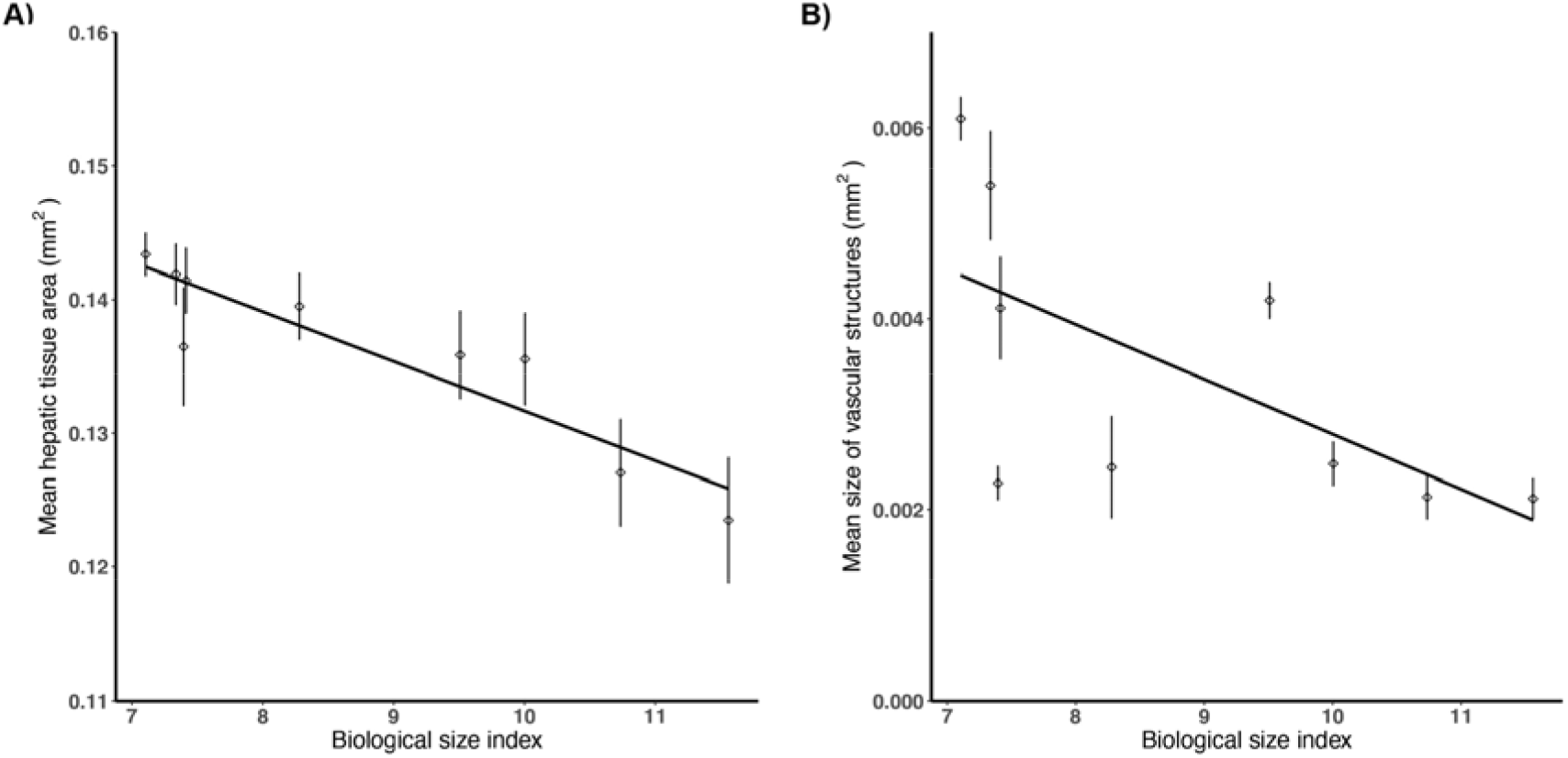
An increase in biological size index (SVL / √ genome size) is correlated with A) a decrease in hepatic tissue area and B) a decrease in the size of vascular structures. Error bars represent standard error.

## Discussion

### Organ Size correlates with body size for the heart and liver

The size of an organism or organ is a function of cell size and cell number. To highlight the importance of this interaction, Hanken and Wake (1993) drew on empirical work in salamanders to introduce the concept of biological size, which is a proxy for the number of cells comprising an organism. We found that ventricle and liver sizes were positively correlated with body size but not genome size in *Plethodon* salamanders, indicating that the size of these organs is a function of body size (Fig. 5, S2–3). Overall, we found no relationship between genome size and body size in *Plethodon*, indicating that increases in cell size do not produce larger body sizes. Thus, the evolution of larger genome and cell size in *Plethodon* is accompanied by a reduction in total cell numbers (i.e., reduced biological size) for both the ventricle and liver. Given this result, we would predict changes in morphology because the organs now develop from and consist of fewer, larger cells.

### Tissue morphology correlates with genome and cell size in the heart and liver

The heart and liver each showed a distinct pattern of phenotypic change accompanying evolutionary increases in genome and cell size in *Plethodon*. In the heart, large genome and cell size was accompanied by a dramatic reduction in the amount of trabeculated myocardia relative to lumen space. Because ventricle size was not correlated with genome and cell size, the reduced myocardial density in species with large genome sizes resulted in hearts with significantly fewer myocardial cells overall.

In the liver, the circulatory structures were most significantly impacted by genome and cell size, which resulted in distinct changes to tissue geometry. The average area of the vascular structures, which included hepatic arteries, portal veins, and sinusoids, was positively correlated with genome size. Conversely, the total number of these vascular structures was negatively correlated with genome size. Thus, livers in species with the largest genome sizes had significantly fewer, but larger, vascular structures. These results suggest that arteries, veins, and liver sinusoids increase in size to accommodate larger blood cells, which in turn changes the geometry and composition of liver tissue. The changes in tissue geometry and composition resulted in two dramatic alterations to overall hepatic morphology. First, the arrangement of vascular structures into a portal triad-like organization was uncommon and became increasingly rare as genome and cell size increased. Although the hepatic arteries, portal veins, and bile ducts were present, they were rarely arranged together in a canonical portal triad. Second, the hepatocytes lacked the 1–2 cell-thick, cord-like morphology, which resulted in numerous hepatocytes having no direct contact with circulating blood. Akiyoshi and Inoue (2012) and others described this same several-cell-thick plate morphology in salamanders (Akat and Göçmen, 2014; Akat and Arkan, 2017; Vaissi et al., 2017). Our results show that this morphology is a result of increased genome and cell size.

The patterns of phenotypic change that we report in the heart and liver, combined with known patterns from brain, skeleton, and circulatory system, suggest that the impact of genome and cell size increase on phenotype is organ-specific (Villalobos et al., 1988; Roth et al., 1993, 1994; Mueller et al., 2008). This lack of similarity suggests that, for each developing organ, unique relationships exist between genome- and cell-level parameters and the morphological outcome of the developmental system. Because large genome and cell size decreases the rate of development, reduces the number of cells involved in morphogenesis, and/or alters the composition and geometry of tissues, a full understanding of the link between cell size and morphology requires understanding which of these effects is relevant for each organ.

### Proposed genome and cell size effects on heart development

We propose three developmental hypotheses explaining the decrease in ventricular musculature associated with increases in genome and cell size. First, this phenotype might arise from an interaction between functional and developmental constraints derived from large cell size. Blood flow plays a significant epigenetic role in vertebrate heart development by creating pressure gradients in the developing heart (Santhanakrishnan and Miller, 2011; Johnson et al., 2015). Large erythrocyte size can potentially change the fluid dynamics of blood, which in turn would change the degree of pressure in the heart during development. The circulation of large erythrocytes has posed other functional challenges resulting in morphological evolution, evident through the repeated evolution of erythrocyte enucleation in miniaturized salamander species (Villalobos et al., 1988; Mueller et al., 2008; Itgen et al., 2019; Decena-Segarra et al., 2020). Testing this hypothesis would require measuring blood flow, its effects on gene expression, and the morphogenetic outcomes in the developing hearts of salamanders with different genome sizes. Second, the reduction of ventricle muscle might be the result of slower rates of development and truncation of heart organogenesis at an earlier ontogenetic stage because of large genome and cell size, producing a paedomorphic heart. Third, morphogenesis and pattern formation can be fundamentally changed when cell size and cell number change (Alberch and Gale, 1985). Because ventricle size was not correlated with cell size, increased cell size must be accompanied by decreased cell numbers. The implications of undergoing heart development with fewer, larger cells are not understood. Testing the latter two hypotheses would require comparative developmental analyses across taxa with different genome and cell sizes to reveal how heart morphogenesis is impacted by changes in developmental rate and in the size and number of cells.

### Proposed genome and cell size effects on liver structure

The most prominent change in liver phenotype as genome and cell size increases is related to tissue geometry and vasculature, which includes arteries, veins, and sinusoids. As genome and cell sizes increase, the sizes of the vascular structures also increase — likely to accommodate larger blood cells — while the number of vascular structures decreases. Thus, large genome and cell sizes are associated with fewer, larger vascular structures. Liver size is determined by body size in *Plethodon* (Fig. 5), which means that increases in cell size are accompanied by decreases in cell number. If the number of vascular structures were unchanged, their increased size would likely cause the tissue to become functionally and/or structurally compromised because they would occupy too much space, at the expense of hepatocytes. Therefore, we hypothesize that the number of vascular structures is constrained to maintain organ functionality and structural integrity.

### Implications of morphology on performance and fitness

The morphological changes that result from increased genome and cell size can have functional consequences. In some cases, functional consequences have been inferred from organs showing patterns of compensatory evolution to offset the negative effects of genome and cell size increase. For example, genome and cell size increase were accompanied by significant changes in brain and eye morphology in salamanders (Roth et al., 1988, 1990). These changes included relative increases in brain and eye size, relative expansion of the regions of the brain responsible for visual and visuomotor functions, increased cell densities within these visual processing brain tissues, optic fibers, and retinas, and a proportional shift in the retina to increase the number of small cones relative to the large rods (Roth et al., 1988, 1990). These compensatory changes in brain and eye morphology offset the low number of large neurons resulting from large cells and small body sizes, maintaining the acuity required for visual predation (Roth et al., 1990).

We hypothesize that heart and liver function are affected by the evolutionary changes we report in *Plethodon* as genome and cell size increase. In the heart, the decreased myocardial volume and increased lumen space in the ventricle likely reduce the ventricle’s capacity to produce force. Conversely, this reduction in myocardium allows the ventricle to hold greater blood volume. A reduction of force production and increased volume capacity would alter the stroke volume and ejection fraction of the heart. Overall functional measures such as heart rate and cardiac output would be affected as well. In the liver, changes in hepatic tissue organization likely affect function as well. Large genome and cell size impacted the organization of portal triads and the cord-like morphology of hepatocytes, as well as reducing the number of vascular structures. As a result, many hepatocytes do not come into direct contact with any vascular structures, limiting their access to oxygen and nutrient supply and thus impacting their metabolic contribution to overall organ function.

It is important to consider, however, whether these cell- and organ-level changes in function have any effect on fitness. Variation in morphology is connected to fitness through its effects on organismal performance (Arnold, 1983). Morphology can vary without impacting performance, and performance, in turn, can vary without impacting fitness (Bock 1980). Are the evolutionary changes that we report here in organ structure, associated with a more-than-doubling of genome size, likely to impact organismal performance and fitness? We argue that the changes likely have little to no effect, as a consequence of another general feature of salamander physiology. Salamanders have incredibly low metabolic rates that appear unrelated to genome and cell size (Gatten et al., 1992; Uyeda et al., 2017; Gardner et al., 2020; Johnson et al., 2021). Low metabolic rates relax selective pressures on the functional capacity of some organs. The heart, for example, is circulating blood in an organism with the lowest O_2_ requirements among terrestrial vertebrates. The low metabolic rates of salamanders also decrease the demand for the liver to metabolize macromolecules (i.e. carbohydrates, lipids, proteins) needed for oxidative phosphorylation and glycolysis. Relaxed selective pressure on organ function allows for greater variation in morphology without negatively impacting organismal performance or fitness. This, in turn, allows cell size to evolve driven largely by genome-level processes (e.g. transposable element proliferation and deletion; Sun et al., 2012). The range of genome and cell sizes produced throughout the clade’s evolutionary history has been funneled through a conserved developmental system, producing a range of morphologies that reflect the crossing of bifurcation boundaries during organogenesis. The “permissive” organismal phenotype of salamanders is thus a powerful tool for examining how the output of developmental systems responds to changes in the fundamental parameter of cell size.

## Acknowledgements

All permits were issued to M.W. Itgen and the animals were collected between May and August of 2018. *Plethodon idahoensis* was collected from Shoshone county, Idaho, under the wildlife collection permit #180226 issued by the Idaho Department of Fish and Game. *Plethodon cinereus* and *P. glutinosus* were collected from South Cherry Valley and Oneonta, Otsego County, New York, under the New York State Department of Environmental Conservation scientific collection permit #2303. *Plethodon vehiculum, P. vandykei*, and *P. dunni* were collected from Pacific County, Washington, under the scientific collection permit # ITGEN 17-309 issued by the Washington Department of Fish and Wildlife. *Plethodon metcalfi* was collected from Macon County, NC, and *P. montanus* and *P. cylindraceus* were collected from Avery County, NC, under the wildlife collection license # 18-SC01250 issued by the North Carolina Wildlife Resources Commission. Specimens of *Ambystoma mexicanum* were obtained from the Ambystoma Genetic Stock Center, which is funded through NIH grant: P40-OD019794. We thank A. Summers, the Summers lab, and the Karel Liem Bioimaging Facility at the University of Washington’s Friday Harbor Laboratories for allowing us access to their CT scanner and for assistance scanning the specimens. For assistance in the field, we thank A. Cicchino, A.H. Griffing, J. Hayes, M. Hayes, E. Itgen, J. Itgen, F. Rodríguez Vásquez, and S.K. Sessions. For discussion of analyses and the manuscript, we thank members of MWI’s dissertation committee K. Hoke, D. Sloan, and W. Zhou. This research was funded by the National Science Foundation (grant 1911585 awarded to RLM), the GREG R.C. Lewontin Early Award awarded to MWI, the Chicago Herpetological Society Grant awarded to MWI, the Stephen and Ruth Wainwright Endowment awarded to MWI, and the Helen T. and Frederick M. Gaige Award awarded to MWI. Animal use was approved by the Institutional Animal Care and Use Committee of Colorado State University and carried out in accordance with protocol 17-7189A.

---

**Table S1.**
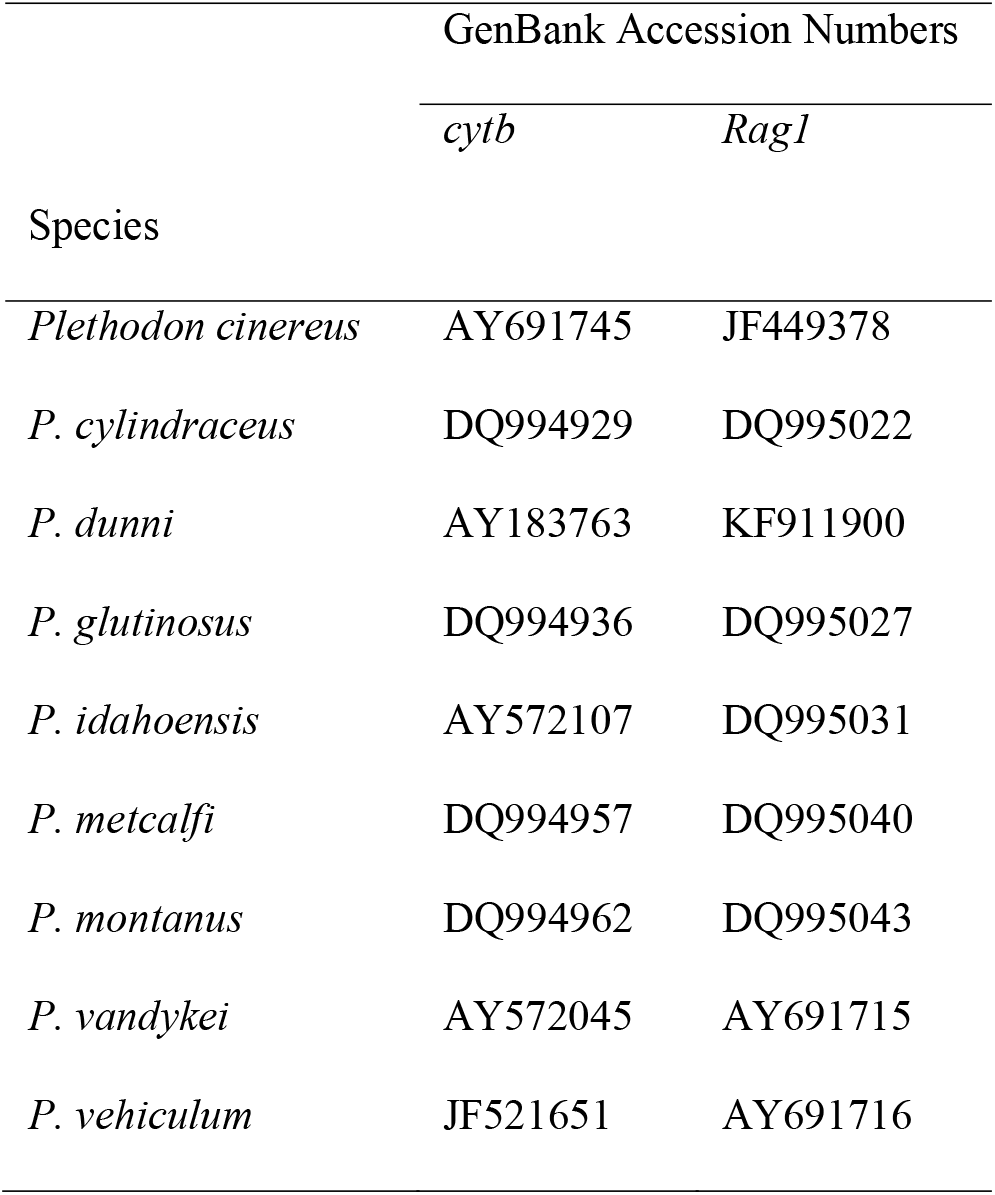
GenBank accession numbers for sequences used in the phylogenetic analysis.

**Table S2.** Estimates of phylogenetic signal for each trait used in the MANOVA analysis using Blomberg’s K. Traits with significant estimates of phylogenetic signal are in bold.

| Traits | Measure of Phylogenetic Signal |  |
| --- | --- | --- |
|  | Blomberg's K | P |
| Snout-vent length (mm) | 0.37 | 0.375 |
| Genome size (Gb) | **1.51** | **0.001** |
| Liver volume (mm<sup>3</sup>) | 0.357 | 0.45 |
| Ventricle volume (mm<sup>3</sup>) | 0.348 | 0.442 |
| Myocardial area (mm<sup>2</sup>) | **0.97** | **0.019** |
| Hepatic tissue area (mm<sup>2</sup>) | 0.564 | 0.136 |
| Number of vascular structures | **2.69** | **0.002** |
| Average vascular structure size (mm<sup>2</sup>) | **2.6** | **0.002** |

**Figure S1.**
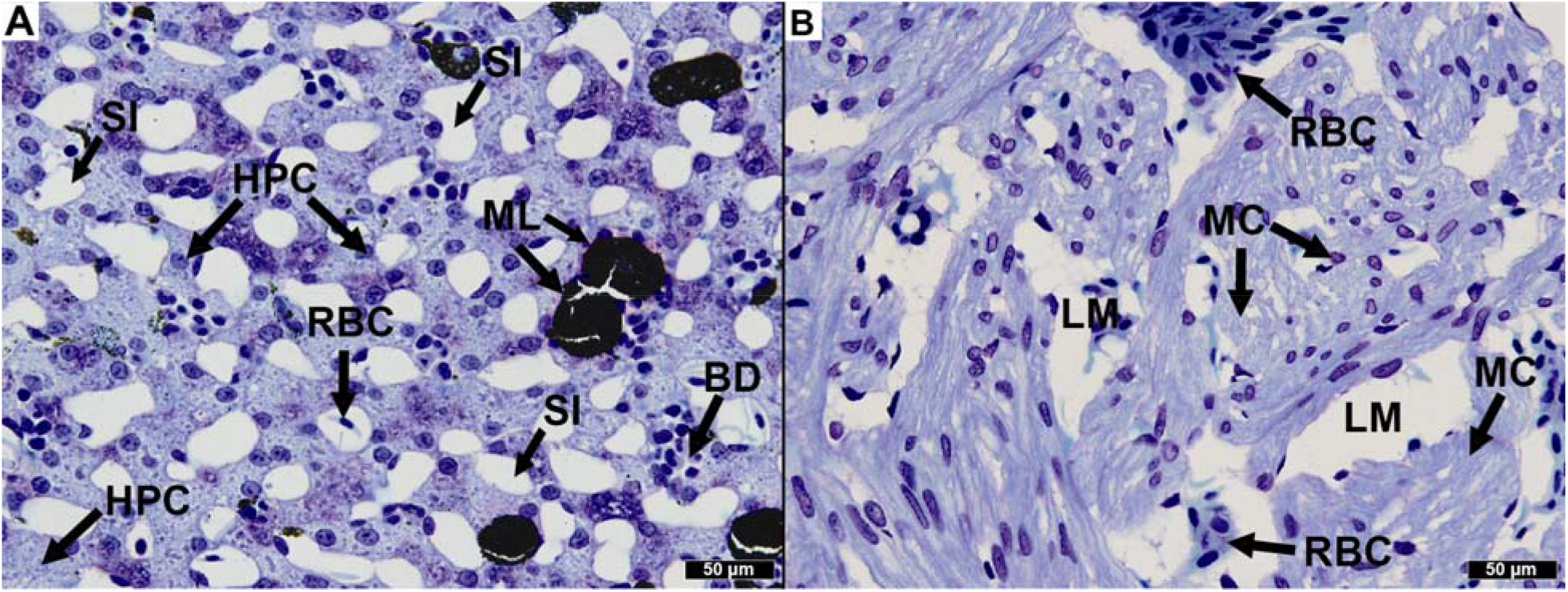
A) Histological section of liver tissue identifying bile ducts (BD), hepatocytes (HPC), melanocytes (ML), red blood cells (RBC), and sinusoids (SI). B) Histological section of the ventricle identifying the lumen (LM), myocardium (MC), and red blood cells (RBC).

**Figure S2.**
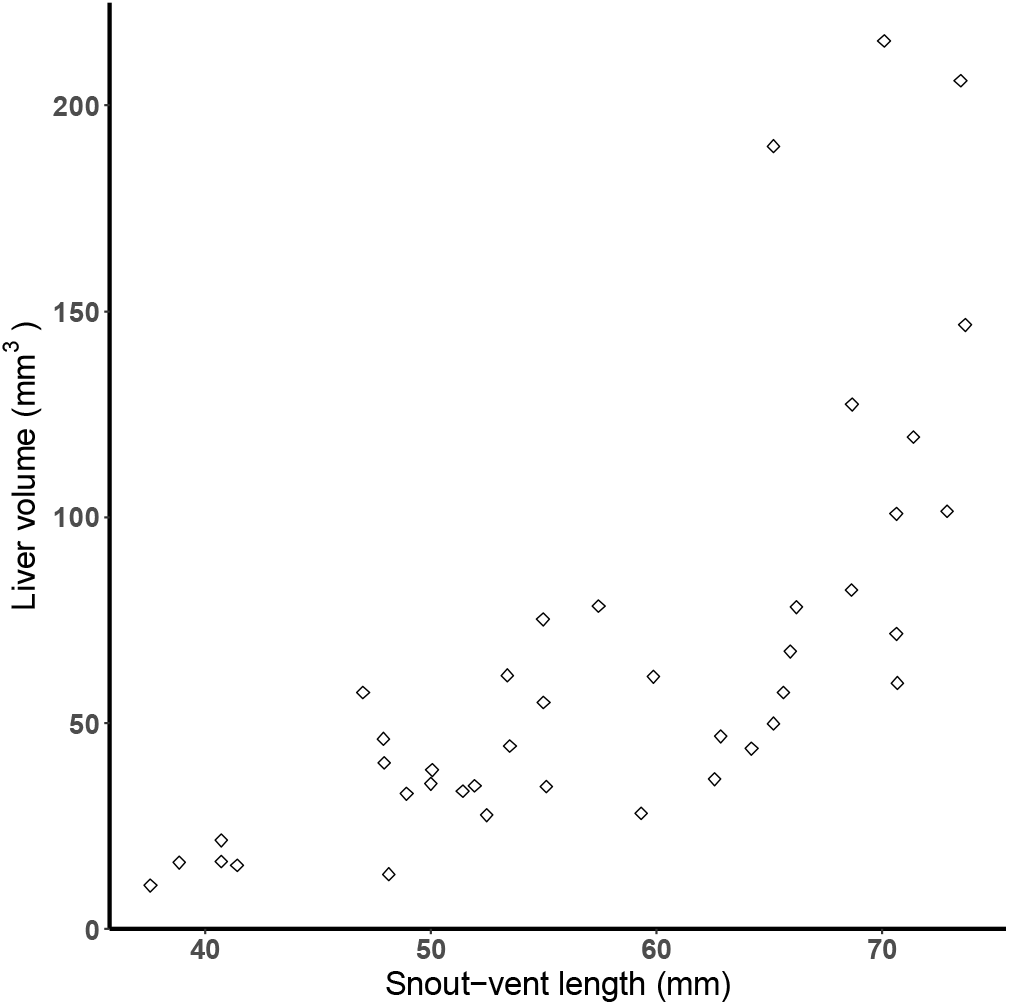
Relationship between liver volume (mm^3^) and SVL (mm) for each individual across all species.

**Figure S3.**
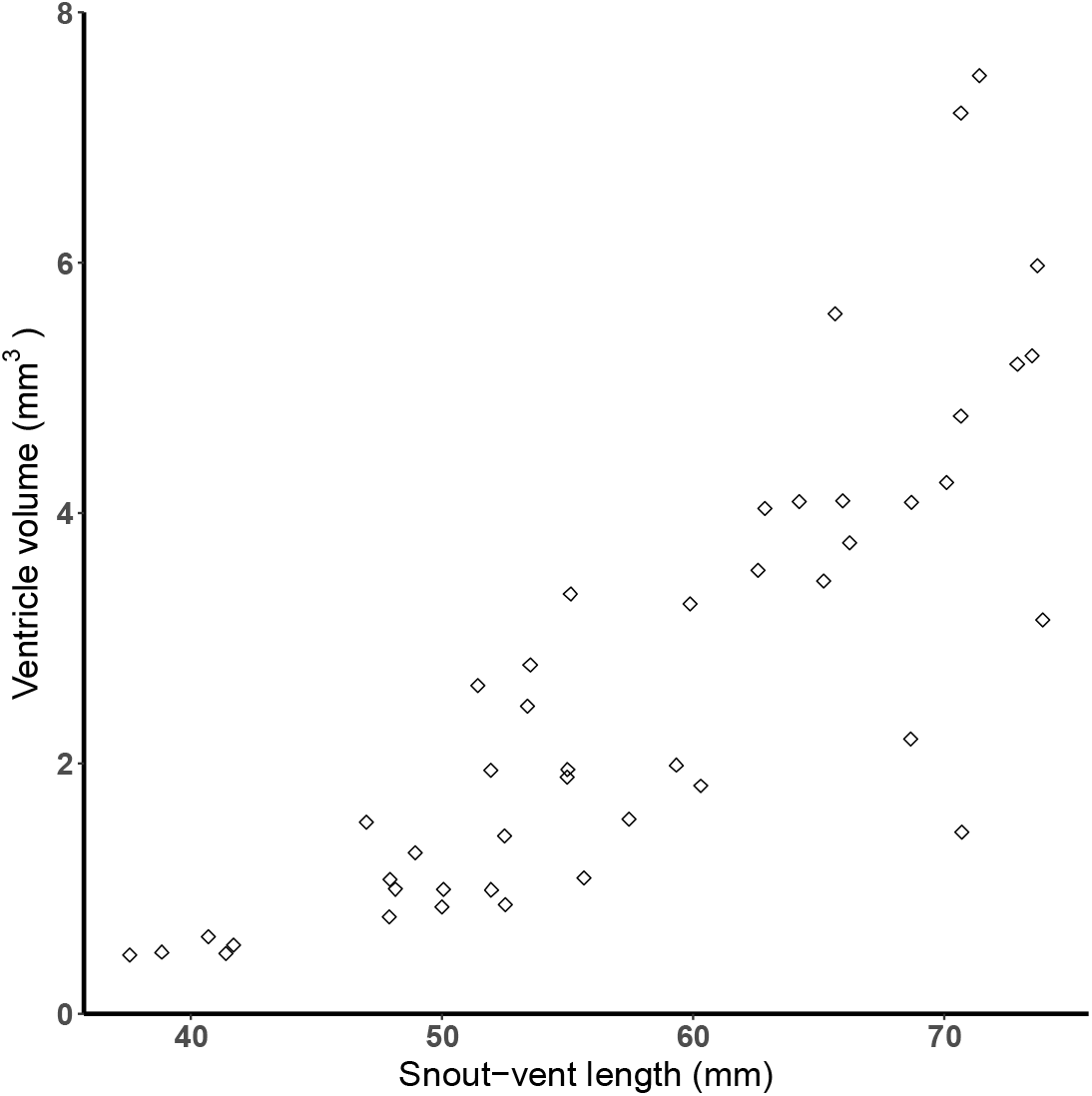
Relationship between ventricle volume (mm^3^) and SVL (mm) for each individual across all species.

